# Sample size dependence of Occam’s razor in human decision-making

**DOI:** 10.64898/2026.01.25.701581

**Authors:** Francesco G. Rinaldi, Eugenio Piasini

## Abstract

To make sense of a noisy world, living beings constantly face decisions between competing interpretations for ambiguous sensory data. This process parallels statistical model selection, where most frameworks, like the Akaike Information Criterion (AIC) and the Bayesian Information Criterion (BIC), are based on a trade-off between a model’s goodness-of-fit and its complexity and prescribe a bias towards simpler models (explanations). This same bias towards simplicity is generally reflected in human behavior. However, a core tenet of normative frameworks is that the trade-off should depend on the sample size (N): as more data becomes available, the goodness-of-fit grows faster than the “cost” of complex models, weakening the overall bias towards simplicity. It is unknown whether humans also conform to an analogous scaling principle, and if so, whether this behavior arises from an internal computation similar to that leading to the normative solution or from a simpler heuristic. Here, we investigate these questions using a preregistered visual task where participants inferred the number of latent Gaussian sources generating clusters of data-points, and where the number of points (N) presented on each trial is varied systematically. We consider three kinds of descriptions for participant behavior: one arising from a linear scaling of the weight of sensory evidence in N (as in BIC and AIC), one with no scaling, and one with sublinear scaling inspired by known biases in numerosity perception. Our results show that the normative, linear scaling description provides the worst account of human behavior. Instead, we find strong evidence for a sublinear scaling of effective sample size. By inferring the shape of this scaling with Gaussian Processes, we reveal two distinct scaling regimes for different ranges of N, consistent with numerosity perception biases. Our findings suggest that, when selecting between competing explanations for sensory data, humans employ an efficient heuristic that repurposes lower-level perceptual mechanisms to dynamically weight evidence against model complexity.

## 1 Introduction

To make sense of the world, living organisms are constantly interpreting sensory data. However, in a world of noisy data and noisier perceptions, more than one interpretation is often possible, with different interpretations leading to completely distinct actions and outcomes. Choosing among competing explanations for sensory data is therefore a crucial task for survival.

When facing tasks that require choosing among different interpretations for some data, humans tend to conform to the principle known as Occam’s razor: the idea that when more than one explanation fits a dataset similarly well, the simpler one is to be preferred [1–10]. For example, when shown some dots sampled by adding noise to an unknown curve, and asked to guess the shape of the curve, participants tended to draw simple, smooth curves rather than squiggly ones [3]. Similarly, when asked to choose between explanations for an unfamiliar disease, humans preferred those with few causes, even if a more complex, multi-cause alternative was significantly more probable [7].

This simplicity bias is, in general, an effective strategy. In fact, Occam’s razor guides most theoretical methods for finding the best model to describe noisy data, a process known in statistics as model selection [11]. In these frameworks, the principle takes shape as a trade-off between maximizing a measure of goodness-of-fit and minimizing penalties for the complexity of the model [12–18]. The particular way in which Occam’s razor emerges in human behavior seems to broadly reflect these normative notions. When asked to choose between two competing interpretations to explain some noisy visual data, participants adopt strategies that reflect the same trade-off, and, moreover, that align with theory-driven notions of complexity [1]. This alignment between human decision-making and the statistical theory of model selection sheds some light on the strategies that humans intuitively employ when faced with these kinds of problems. It is still unclear, however, if the alignment stems from the application of similar theoretical principles, perhaps implemented as a probabilistic computation in the brain, or if humans simply employ heuristics that mimic the theoretical definitions of complexity while adopting a more elementary approach.

In theoretical frameworks the simplicity bias emerges from the application of optimality principles. For example, minimizing the distance between the unknown true generating distribution *g*(*x*) for some data *X* and a parametric distribution *f*(*x* | *θ*_*M*_) from a candidate model *M* results in the well-known Akaike Information Criterion (AIC): *M*_*best*_ = max_*M*_ {*L*_*M*_ − *K*_*M*_} where the maximum log-likelihood of the model, *L*_*M*_, is the goodness-of-fit measure, and its number of parameters, *K*_*M*_, is the complexity penalty [12]. Similarly, in the Bayesian model selection (BMS) framework the optimal model is the one with the highest posterior probability given the observed data: 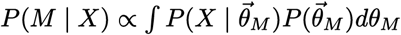, where 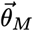 are the model parameters [17]. Even though BMS does not explicitly introduce any measure of complexity, for large N the logarithm of the integral can be approximated with an asymptotic expansion as a sum of terms, each growing more slowly than the previous as N increases. Keeping only the first two terms of the expansion, one obtains the Bayesian Information Criterion (BIC): log *P*(*M* | *X*) ≃ *L*_*M*_ − *D*_*M*_ + *O*(1), where we have again the log-likelihood term and a penalty know as Dimensionality, given by *D*_*M*_ = (log(*N*)/2)*K*_*M*_, with *N* number of data-points (Fig. 1a) [13]. Continuing the expansion produces more penalty terms, corresponding to more nuanced characterizations of model complexity that can be interpreted with the tools of information geometry [1, 18]. Analogous notions of complexity can be derived with different approaches, including the Minimum Description Length principle [19, 20] and Predictive Information [21].

**Figure 1.**
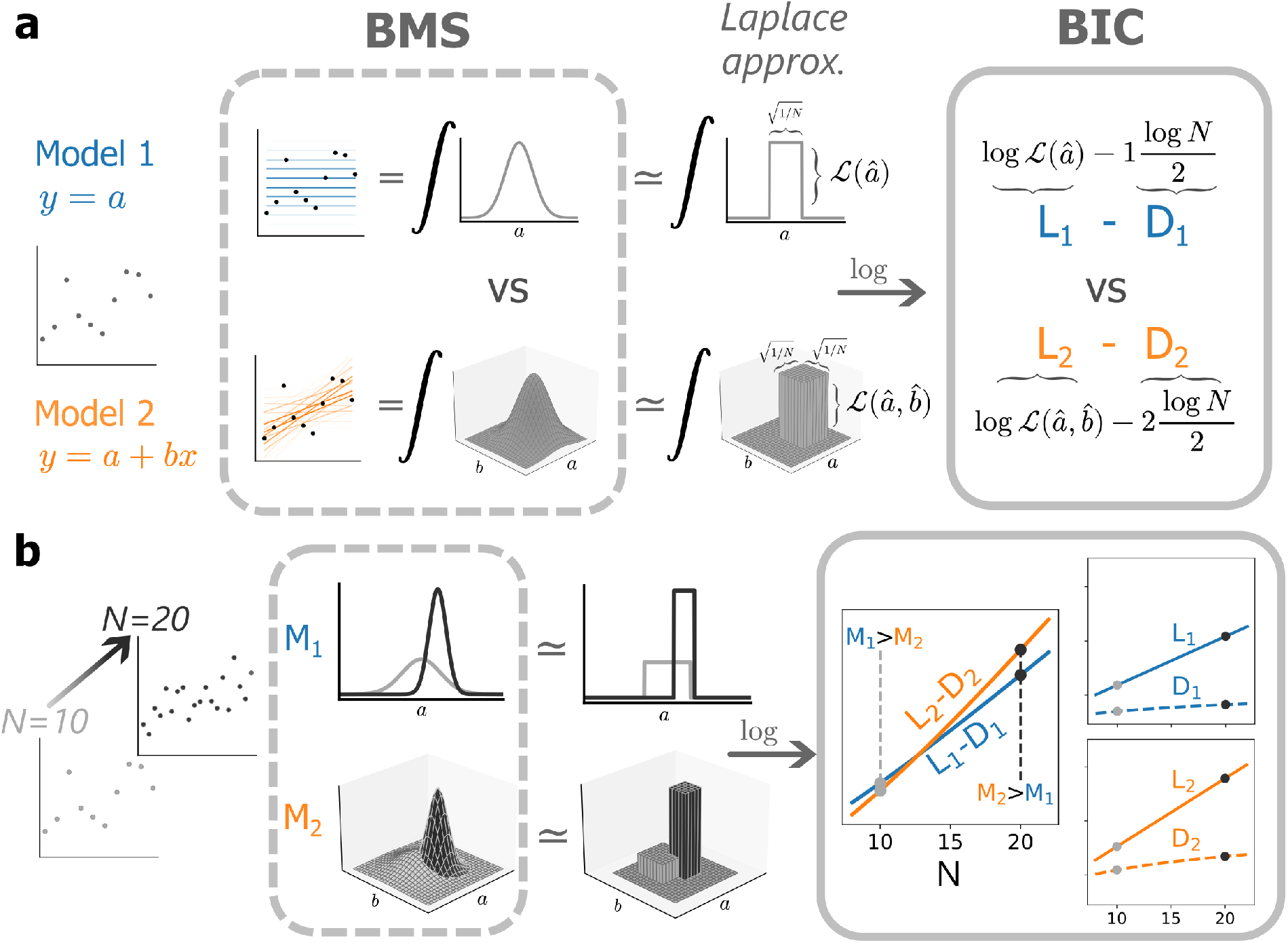
Bayesian Model Selection balances goodness-of-fit and complexity of statistical models. **a**. Bayesian Model Selection (BMS) between a model with one parameter (Model 1, linear model with fixed slope = 0, blue) and a model with two parameters (Model 2, linear model, orange). Given the same prior for both models and a uniform prior across their parameters, the best model will be the one with the highest likelihood averaged across all the possible choices of parameters (dashed box, left). This quantity, the marginal likelihood, corresponds to the integral of the likelihood times the prior (dashed box, right). By using the Laplace approximation, this integral can be decomposed into the product of a “height” term (proportional to the maximum likelihood of the model) and “width” terms (proportional to the inverse square root of the number of data-points *N*). Taking a logarithm, these terms correspond to the log-likelihood *L*_*M*_ and the dimensionality *D*_*M*_ of the Bayesian Information Criterion (BIC, solid-edge box). **b**. Effect of increasing the sample size. The height of the likelihood landscape is exponential in the number of points *N* (since it is the product of the single-point likelihood of *N* data-points), while the widths of the peaks decrease as *N*^−1*/*2^. Taking a logarithm, this translates to a log-likelihood term that grows linearly in *N* and a dimensionality term that grows logarithmically. Since the goodness-of-fit grows faster than the complexity penalty, the simplicity bias gets relatively weaker for higher sample sizes, allowing for the selection of more complex models (solid-edge box).

In all these frameworks, the strength of the simplicity bias changes when changing the amount of data available for the inference. For independently drawn observations, the likelihood is the product of the probability of each data-point, and thus the log-likelihood will be the sum of their log-probabilities. *L*_*M*_, therefore, grows on average linearly with the number of observations *N*. On the other hand, the complexity penalties are either independent or sublinearly dependent on the quantity of observed data. The result is that the importance of complexity relative to goodness-of-fit becomes smaller as the amount of available data increases. This is a major component in every implementation of Occam’s razor: when a larger sample is available, a rational decision-maker can afford the use of more complex models (Fig. 1b).

As we have seen, previous work has established the presence of a simplicity bias in humans and quantified its strength. However, it is not known how this strength and its importance relative to goodness-of-fit depend on the amount of data that is available to human observers, as the works that touched upon the matter did not attempt to characterize the simplicity bias quantitatively [10, 22]. Moreover, if such a scaling exists, it is not known if it is a side effect of an approximate computation, such as Bayesian integration over unknown causes mirroring closely the emergence of Occam’s razor in the normative frameworks discussed above, or if it emerges from an explicit heuristic mandating that complexity should become less and less important for larger sample sizes. In the latter case, one would expect that the scaling would be affected by cognitive processes and distortions related to numerosity estimation [23–26].

Here we investigate the existence and nature of this scaling. To do so, we focus on AIC and BIC as normative frameworks to describe human behavior in a visual model selection task. The complexity penalties of these frameworks, *K*_*M*_ and *D*_*M*_ respectively, depend in a simple and well understood way on the sample size *N* (Fig. 1). Moreover, the dimensionality *D*_*M*_ was shown to be the most important term in shaping human strategies, making it an ideal factor for this study [1]. If humans intuitively perform a computation based on optimality principles (for instance, if they perform an approximate Bayesian integration), we expect their behavior to be best described by a linearly growing goodness-of-fit counterbalancing a constant (*K*_*M*_) or sublinearly growing (*D*_*M*_) penalty, as in the normative frameworks. If instead there is no scaling in N, and human choices are only guided by an empirical estimate of the true distribution of sampled data, we expect the best description to have a fixed penalty and a goodness-of-fit independent of *N*, such as the divergence between the empirical measure of the data and a candidate model distribution. Finally, if human strategies are based on a heuristic that explicitly requires scaling based on the estimated number of data-points, we would expect the growth of both the goodness-of-fit and the penalty to be distorted compared to their theoretical counterparts, consistently with the well-known sublinear effects that can be observed in human numerosity perception [24, 26–28].

Our results show that human behavior is best described by a complexity penalty whose importance relative to the goodness-of-fit grows sublinearly as the sample size N grows. By using Gaussian Processes to estimate the precise shape of the scaling, we show that it matches a logarithmic trend, consistently with the prescription of Weber’s law [27], up to high values (≃ 50) of *N*, where a transition happens. This trend could match the transition observed in visual numerosity perception between a regime of direct number estimation and a regime of estimation from density [26, 28], reinforcing the idea that the scaling stems from a heuristic-mandated weighting based on estimated sample size.

## 2 Results

We designed a perceptual decision-making task to investigate the strategies of intuitive model selection employed by human participants. Participants were shown noisy visual data and had to guess which one among a set of possible models generated it. To test a wide range of dimensionalities *D*_*M*_, we used a set of 5 models, each with a different number of parameters. Each model was a mixture of *K* one-dimensional Gaussian components, with *K* ∈ [1, 5] (Fig 2a, bottom). The standard deviation of all components was fixed (*σ* = 0.1), and within a model all components had the same weight *π* = 1/*K*, thus limiting the parameters of the model to the *K* means of its components. These constraints allowed us to make the task more manageable for the participants, as well as focus on a single kind of parameter, avoiding the possibility that different kinds of parameters might have a different influence on human strategies.

**Figure 2.**
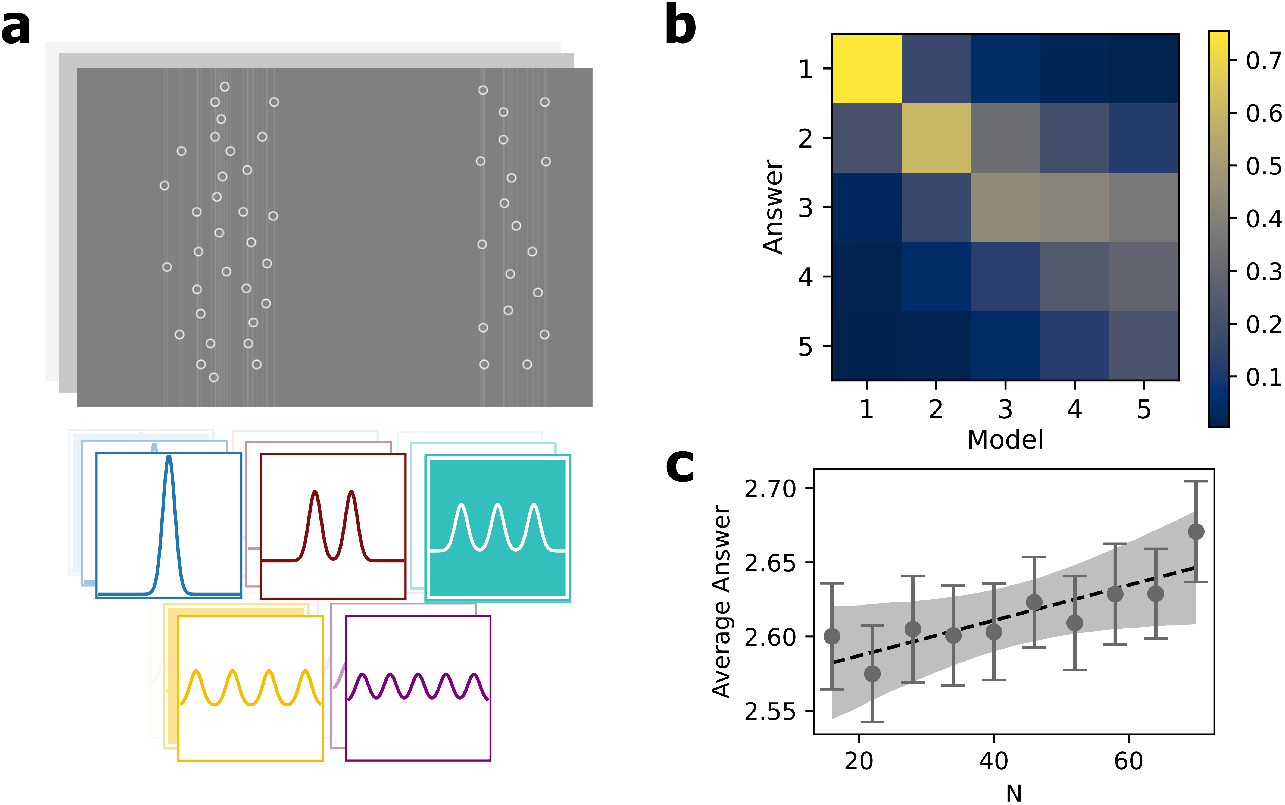
Task description and summary of participants’ answers. **a.** The five models that participants had to choose among (bottom) and an example of a trial from the task (top). The trial shown was generated by a model with three components; two had close means and generated the cluster on the left, one had an isolated mean and generated the cluster on the right. **b**. Average confusion matrix across participants and trials: while for models with low dimensionality (*K*_*M*_ = 1, 2) the answers mostly correspond to the true model, when the number of parameters increases, the answers are skewed towards values lower than the true ones, showing the effect of the simplicity bias. **c**. Average answer across all participants and trials as a function of the number of data-points *N*. Gray markers denote the mean across participants, with error bars representing the standard error. The dashed black line represents the posterior mean of a Bayesian linear regression on the individual participants’ mean responses for each *N* (slope = (1.2 ± 0.6) × 10^−3^, prob. of direction = 0.026). The shaded gray region denotes the 95% Highest Density Interval (HDI) of the expected value. The overall value of the average answer remains below the actual average (⟨*K*_*M*_⟩ = 3), confirming a persistent simplicity bias. However, the positive slope captured by the inference suggests a gradual weakening of this bias as the amount of available data (*N*) increases.

On each trial, data were generated by one of the models, selected at random with uniform probability. The positions of the *K* means for the selected model were randomly sampled on a horizontal line [−1, 1], again with uniform probability, and *N* observations were sampled from the resulting mixture distribution. These observations were represented as white vertical lines on a gray background. To avoid underestimation of the sample size due to overlapping lines, each line was marked by a white circle (Fig 2a, top). To study the effect of the amount of available data we showed trials with ten different sample sizes *N*, equally spaced between 16 and 70. The amount of trials generated by each model for each value of *N* was balanced to ensure a uniform exposure to all the models. Overall, the experiment consisted of 900 trials, shown in blocks of 30. Within a block, all trials had the same sample size. We collected data from 100 participants (see Methods for more details).

An initial analysis of the data revealed that, on average, task performance was lower on trials where the true value of *K* was higher, and participants’ error patterns were skewed towards reporting a value of *K* lower than the true one (Figure 2b). Moreover, the average value of *K* reported by participants increased as a function of *N* (Figure 2c). Overall, these patterns suggest that participants exhibit a simplicity bias (preferring models with lower *K*), and that this bias becomes weaker as the amount of available data increases.

To test these hypotheses further, we turned to a statistical analysis pipeline that we designed and preregistered before collecting the data [29]. The pipeline uses a Bayesian multinomial logistic regression [30] to measure the degree to which each participant’s choices were affected by the goodness-of-fit and the complexity penalty of models [1]. Specifically, we assigned a measure of sensitivity *β* to each of the terms: the probability to choose a specific model was therefore dependent on the sum of the product of each term (goodness-of-fit or penalty) with its sensitivity parameter:

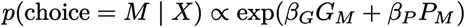

Where *G*_*M*_ is the goodness-of-fit of the model *M* for the data *X* and *P*_*M*_ is its complexity penalty.

We first verified whether the behavior of participants displays a simplicity bias in our setup. We compared regressions based on AIC-like or BIC-like decision rules (*G*_*M*_ = *L*_*M*_ and respectively *P*_*M*_ = *K*_*M*_, the number of parameters, or *P*_*M*_ = *D*_*M*_, the dimensionality) with a regression that only uses the log-likelihood *L*_*M*_, without any penalties. The comparison was performed using Pareto smoothed importance sampling Leave-One-Out Cross-Validation (PSIS-LOO CV, [31]). This cross-validation method allows us to approximate the Expected Log Predictive Density (ELPD) of a fitted regression, a statistical measure of its predictive power on unseen data [32]. As shown in Fig. 3a, AIC and BIC greatly outperform the predictive power of a likelihood-only regression. This confirms that our participants penalize complex models, following Occam’s razor, while performing the task.

**Figure 3.**
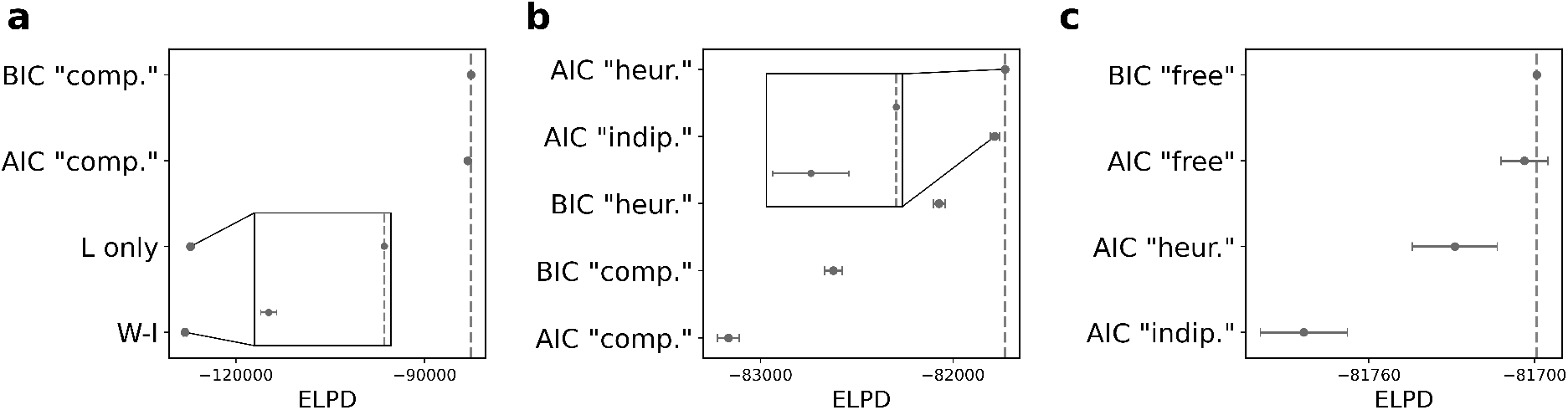
Model comparison between candidate descriptions. **a**. Pareto-Smoothed-Important-Sampling Leave-One-Out Cross-Validation (PSIS LOO CV) comparison between the two Goodness-of-fit descriptions (W-I and Likelihood-only) and two descriptions based on AIC and BIC. The x-axis shows the Expected Log Predictive Density (ELPD) of each description. The error bars represent the standard error of the ELPD differences with the best model, ELPD_**best**_-ELPD_*M*_. The inset shows the PSIS LOO comparison between the Likelihood-only and the W-I descriptions, showing that the Likelihood-only description has a significantly higher ELPD than the W-I one. **b**. ELPD comparison between the “computation”, “independent” and “heuristic scaling” description. The “computation” ones perform significantly worse than the others. The inset highlights that the difference between the AIC “heuristic” and AIC “independent” descriptions is within three standard deviations. **c**. ELPD comparison between the AIC “heuristic”, AIC “independent” and “free scaling” descriptions. The “independent” description is the only one to perform significantly worse than the others.

Moreover, we found that the inferred sensitivity to dimensionality *D*_*M*_ in the BIC-like description (and similarly the sensitivity to *K*_*M*_ in the AIC-like one) is greater than the sensitivity to likelihood, as well as the theoretically optimal value of 1 (See Tables 1 and 2). This over-weighting of dimensionality has also been observed by [1]^1^.

**Table 1:** Posterior summary statistics for the key population-level parameters of the AIC “computation” description. The columns represent the posterior mean (**mean**), posterior standard deviation (**sd**), the 3% and 97% boundaries of the Highest Density Interval (**hdi_3%, hdi_97%**), as well as a number of standard diagnostics for MCMC-based Bayesian inference, namely the Monte Carlo Standard Error for the mean and standard deviation (**mcse_mean, mcse_sd**) [30], the bulk and tail Effective Sample Size (**ess_bulk, ess_tail**) [56], and the R-hat convergence diagnostic (**r_hat**) [56]. All diagnostics were computed with ArviZ version 0.15.1 [57].

|  | mean | sd | hdi_3% | hdi_97% | mcse_mean | mcse_sd | ess_bulk | ess_tail | r_hat |
| --- | --- | --- | --- | --- | --- | --- | --- | --- | --- |
| $\mu_{\beta_L}$ | 0.232 | 0.016 | 0.200 | 0.258 | 0.000 | 0.000 | 3886.6 | 3857.8 | 1.000 |
| $\mu_{\beta_D}$ | 1.423 | 0.059 | 1.308 | 1.531 | 0.001 | 0.001 | 3945.6 | 3795.4 | 1.001 |
| $\sigma_{\beta_L}$ | 0.152 | 0.012 | 0.130 | 0.175 | 0.000 | 0.000 | 3908.6 | 3774.8 | 1.001 |
| $\sigma_{\beta_D}$ | 0.598 | 0.045 | 0.514 | 0.679 | 0.001 | 0.000 | 4096.5 | 4013.6 | 1.000 |
| $\mu_{\epsilon}$ | 0.023 | 0.003 | 0.018 | 0.028 | 0.000 | 0.000 | 2931.7 | 3497.9 | 1.001 |

**Table 2:** Posterior summary statistics for the key population-level parameters of the BIC “computation” description. Details are as in Table 1.

|  | mean | sd | hdi_3% | hdi_97% | mcse_mean | mcse_sd | ess_bulk | ess_tail | r_hat |
| --- | --- | --- | --- | --- | --- | --- | --- | --- | --- |
| $\mu_{\beta_L}$ | 0.236 | 0.016 | 0.205 | 0.265 | 0.000 | 0.000 | 4390.1 | 4080.2 | 1.000 |
| $\mu_{\beta_D}$ | 0.798 | 0.035 | 0.733 | 0.867 | 0.001 | 0.000 | 4054.9 | 3553.4 | 1.001 |
| $\sigma_{\beta_L}$ | 0.156 | 0.012 | 0.134 | 0.179 | 0.000 | 0.000 | 4022.6 | 3927.5 | 1.000 |
| $\sigma_{\beta_D}$ | 0.343 | 0.026 | 0.295 | 0.390 | 0.000 | 0.000 | 3842.7 | 3962.7 | 1.000 |
| $\mu_\epsilon$ | 0.023 | 0.003 | 0.018 | 0.028 | 0.000 | 0.000 | 3041.1 | 3563.4 | 1.000 |

We then tested whether a more elementary, intuitive measure of goodness-of-fit would better align with human strategies. We compared the regressions described above with a decision rule based on a set of heuristics tailored to the nature of the sensory stimuli. Since the models that our participants compare generate clusters of points, we used as features the cluster width (W) and the imbalance of points between clusters (I): a good model would be able to partition the data into similarly wide and similarly populated clusters. Each term had its own sensitivity parameter *β*, allowing the regression to flexibly find the right weights for the two goodness-of-fit metrics. The ELPD of the resulting regression (W-I regression) was lower not only than the ones incorporating a complexity penalty (Fig. 3a), but also of the likelihood-only inference (Fig. 3a, inset), showing that, compared to these heuristic features, *L*_*M*_ captures more of the information used by participants to drive their choices.

We therefore proceeded to study whether and how goodness-of-fit and complexity scale with sample size. We developed a total of five regression variants for AIC and BIC-based terms. The variants are divided into three kinds (“computation”, “independent”, and “heuristic scaling”), one for each possible behavior of the scaling (Fig. 4). The two “computation” variants (AIC_comp_ and BIC_comp_) represent decision strategies based on a direct computation, similar to the ones that lead to the AIC and BIC criteria and that could, for instance, be performed through probabilistic computations implemented at the neuronal level [33, 34]. These variants make use of the theoretical values for the goodness-of-fit and the complexity penalty (Fig. 4, left). These are the variants used in the previous comparison. The “independent” variant (AIC_ind_) matches a strategy that doesn’t depend on the number of points, but only on their density. Both the goodness-of-fit and the penalty are independent of the sample size. To implement it, we substitute the log-likelihood *L*_*M*_ with the average point-wise log-likelihood *l*_*M*_ = *L*_*M*_ /*N* (Fig. 4, center). This quantity corresponds to the Kullback-Leibler divergence between the empirical measure of the data and the candidate model distribution, up to an additive term (the entropy of the empirical measure of the data, which doesn’t change between models and thus doesn’t affect the probability *P*(choice = *M* | *X*)). In this case, we only consider one variant in common for AIC and BIC, since, if we remove the dependence on *N* of the BIC dimensionality *D*_*M*_, it will be the same as the AIC penalty *K*_*M*_ up to a multiplicative factor. Finally, to check whether cognitive distortions affect human strategies, in the two “heuristic scaling” variants (AIC_heur_ and BIC_heur_) we substitute the dependence on *N* in each term with a dependence on *Ñ* = log *N*. This compressed dependence represents the effect of a sublinear Weber-like perception of numerosity [24]. Therefore, the goodness-of-fit term will be *Ñl*_*M*_ and the BIC dimensionality penalty log(*Ñ*)*K*_*M*_ /2, while the AIC penalty remains unchanged (Fig. 4, right). The details of all these variants, with the exception of the “independent” one, were specified before data collection in our preregistration document [29].

**Figure 4.**
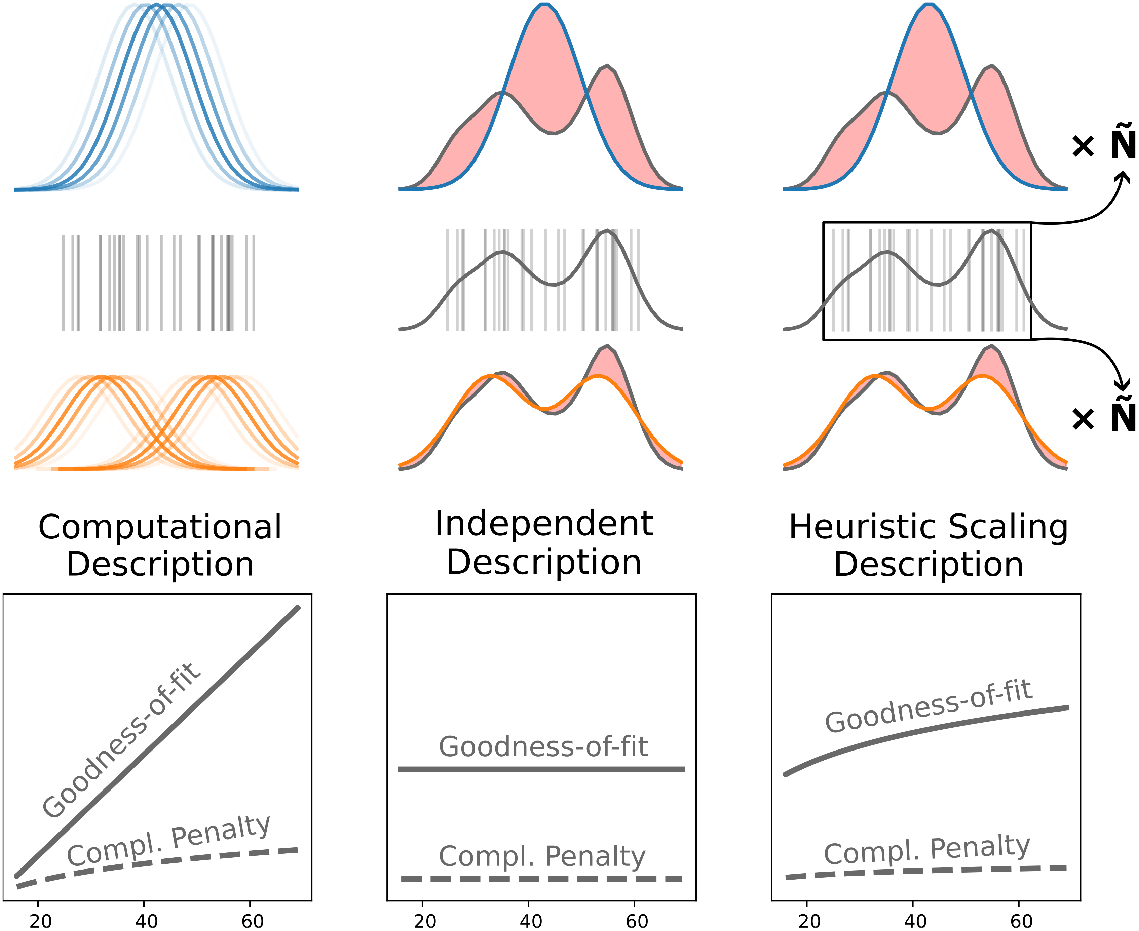
Schematic illustration of the three descriptions we used to model the participants’ strategies and dependence of their terms on the sample size *N*. Only the BIC versions are shown. The “computation” description (top left) assumes that participants’ choices depend on a computation similar to Bayesian integration: as shown in Fig 1a, the integration sums the likelihoods for each possible parameter value, here represented as different values of the components’ means. The resulting goodness-of-fit scales linearly in *N*, while the resulting complexity penalty scales logarithmically (bottom left). The “independent” description (top center) assumes that participants only rely on an estimate of the density of the data (gray line) and compare it with the best-fitting shape of the models. In the figure, the red areas represent the difference between the empirical data distribution and the model distributions. Both the goodness-of-fit and the dimensionality are constant in *N* (bottom center). The “heuristic scaling” description (top right) assumes that participants rely on a process similar to that in the “independent” description, but try to recover the optimal scaling by weighting the result by an “effective” perceived number of data-points *Ñ*. The resulting goodness-of-fit scales as the effective numerosity of the data, which we assume to be *Ñ* = log(*N*) [27], while the penalty scales as *Ñ*.

We used LOO-CV again to perform the comparison (Fig 3b). The “computation” variants are outperformed by all other variants. This suggests that in our task participants don’t closely follow a decision rule that is well described by theory-like computations, such as (approximate) Bayesian integration. However, it is harder to adjudicate between the other hypotheses and to draw a definite conclusion on the scaling of the simplicity bias with the sample size *N*. The AIC “heuristic” variant, where the importance of goodness-of-fit scales as *Ñ* = log(*N*), is the one with the highest predictive power, but it is closely followed by the AIC “independent” variant, where there is no scaling at all (Δ*ELPD* = 54.65 ± 24.38; Fig 3b, inset).

To better understand the scaling trend, we therefore developed two more variants for AIC and BIC, the “free scaling” variants (*AIC*_free_ and *BIC*_free_). These variants are operationally the same as the “heuristic scaling” variants, but instead of substituting the dependence on *N* of the terms with a dependence on *Ñ* = log *N*, we use a generic function *Ñ* = *f*(*N*). The function is implemented as a Gaussian Process (GP) [35], inferred together with the other parameters of the regression. This allows us to give a flexible estimate of the shape of the scaling, and thus to distinguish between a flat dependence (as in the “independent” variant) or a sublinear one (as in the “heuristic scaling” variant). The inferred curves for the “free scaling” variants are shown in Fig 5a. They both follow a sublinear trend up to *N* ≃ 50, where the *Ñ* of AIC_free_ flattens and the *Ñ* of BIC_free_ starts to decrease. These trends suggest that participants follow a “heuristic scaling” type of behavior up to high values of *N*, where the strategy seems to change.

**Figure 5.**
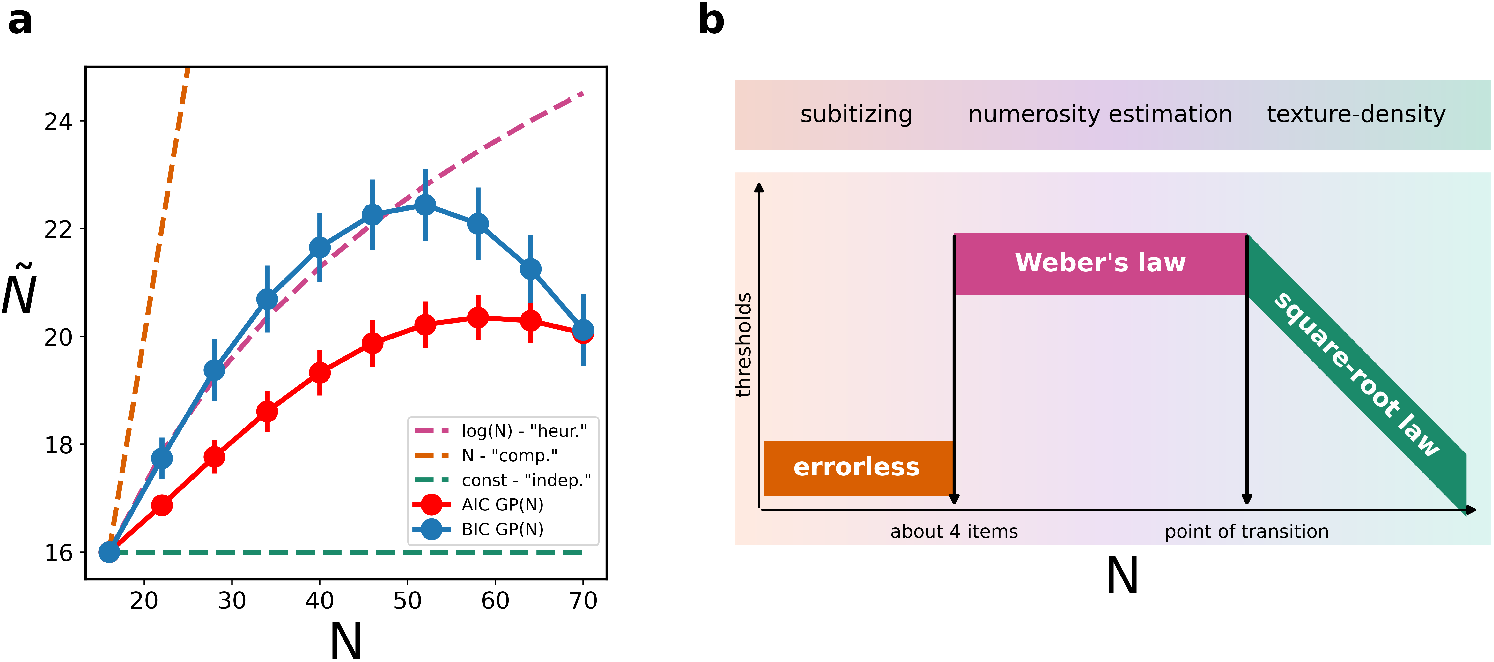
Scaling of the effective sample size. **a**. The inferred functions *Ñ* = *GP*(*N*), computed for both the AIC (red) and BIC (blue) “free scaling” descriptions, compared with the linear scaling of the “computation” descriptions (gray), the flat scaling of the “independent” description (green) and the logarithmic scaling of the “heuristic scaling” descriptions (pink). Both the AIC and the BIC functions follow a sublinear trend for low sample sizes, with a prominent transition around *N* = 50 where *Ñ* starts to flatten out or decrease. This transition may highlight (and compensate) a shift in the underlying heuristic applied by participants happening around this value of *Ñ*. **b**. The three stages of numerosity perception: errorless (orange), Weber-like (pink), and density-based (green). Image adapted from [26].

A possible explanation for this transition, at least for the *Ñ* of AIC_free_, is a strategy change from a numerosity-based heuristic to a density-based one, and therefore from a strategy dependent on the number of data-points to an independent one. This kind of change is reminiscent of the observed switch from Weber-like numerosity perception to density-based estimation for large numbers of data-points (Fig 5b, [26, 28]). However, in the literature, the specific value (or range of values) of *N* where this switch occurs is reported to depend on the details of the experimental setup: in [28], the transition threshold depended on the eccentricity of the cluster of data-points or on the distance between clusters. In our setup, not only do we not control for the distance of the stimulus from the center of vision, as the participants are free to move their gaze, but different trials display clusters in different positions, and often more than two clusters are presented at once, making it difficult to link the two perceptual phenomena in a more quantitative way. Moreover, our analysis doesn’t allow us to infer the Weber fractions and compare them to the literature, as their value is impossible to disentangle from that of the sensitivity to goodness-of-fit *β*_*L*_.

Finally, while LOO-CV allowed us to identify the “free scaling” descriptions as those with the highest relative predictive power (Fig 3c), it does not provide an absolute measure of how well they capture human behavior. To evaluate this, we compute the absolute predictive accuracy of BIC_free_, the best performing description. We defined a *description consistency* as the average fraction of trials where the description’s highest-probability answer coincided with the participant’s actual choice. We compared this to a *participant self-consistency* measure, similarly defined (exploiting the fact that each trial is repeated three times throughout the task) as the average fraction of trials where the most common answer across a triplet of repetitions coincided with the participant’s choice in each repetition. This within-participant reliability measure serves as an upper bound for the description consistency. As shown in Fig 6, the description consistency closely approaches the participant’s self-consistency. By using random chance as the lower baseline and the participant’s self-consistency as the highest one, we obtain a scale of relative description performance (Fig S1), showing that BIC_free_ captures an average of 71% of the explainable behavioral consistency.

**Figure 6.**
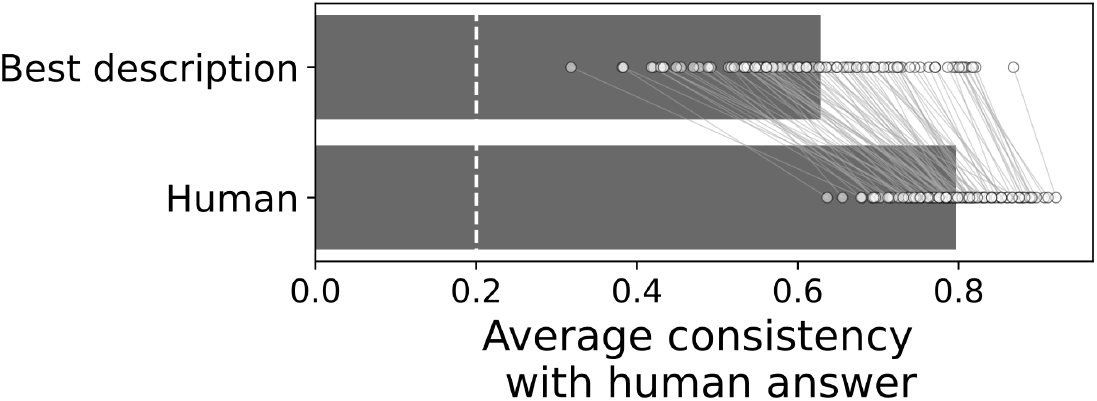
Absolute predictive accuracy and within-participant reliability. The horizontal bars represent the average consistency across all participants. The “Human” bar (self-consistency) shows the average fraction of trials where the participant’s choice in a given repetition matched their most common choice across a triplet of identical trials. The “Best description” bar (description consistency) shows the average fraction of trials where the highest-probability answer of the BIC_free_ description matched the participant’s actual choice. The scattered dots represent the consistency of individual participants, with thin lines connecting the self-consistency and description consistency for the same participant. The vertical dashed line indicates the consistency that would be achieved by randomly guessing the participant’s choices (chance level of 0.2). The reported consistency corresponds to an average normalized consistency between model and participants of 71%, see Figure SS1.

## 3 Discussion

This study investigated how sample size influences human decision-making when choosing an explanation for noisy sensory data. In particular, we studied how the amount of available data influences the trade-off between favoring well-fitting explanations and penalizing their complexity. While previous research has established that humans have a bias towards simple explanations, aligning with the principles of Occam’s razor, it remained unknown whether this bias adapts to the sample size (*N*) as prescribed by normative statistical theories [12, 13]. Our results confirm that, as previously shown, this simplicity bias can be quantified using experiments inspired by the theory of Bayesian model selection [1]. Moreover, we demonstrate that human choice strategies do change with the number of data-points, but without following the linear scaling of the relative importance of the goodness-of-fit that would be expected from an approximate Bayesian computation or other optimal statistical methods. Instead, our results strongly support the existence of a sublinear scaling, where the influence of goodness-of-fit grows more slowly than the amount of data. The shape of this sublinear scaling, which seems to follow different trends depending on the sample size (logarithmic for smaller sample sizes and flattening or decreasing for larger ones) hints at the existence of composite heuristics, paralleling the cognitive phenomena observed in numerosity perception [26, 28]. This suggests that an estimate of the quantity of data-points might be used by participants to inform their balance between complexity and goodness-of-fit.

Inference frameworks such as Bayesian Model Selection provide a principled reason for why the penalty for complexity should become less important as more data becomes available [18, 36]. The maximum log-likelihood, representing goodness-of-fit, grows linearly in *N* for independent data-points, while complexity penalties grow much slower (as in BIC) or not at all (as in AIC). This change of balance naturally arises from the application of optimality principles and ensures that with enough evidence, a more complex model can be justifiably chosen. The mismatch between this normative prescription and what we observe in our data suggests that participants implement a fundamentally different, and potentially more resource-rational, computational strategy.

Our results are consistent with strategies that don’t compute goodness-of-fit through an explicit calculation of the maximum likelihood, by summing the log-likelihood of individual points. Human behavior is instead better described by a process that incorporates a global evaluation of goodness-of-fit for the whole dataset. This evaluation could, for instance, be based on the difference between the data density and the model shape, which could make it more computationally efficient. At the same time, our participants implicitly assigned a weight, or a measure of trustworthiness, to this quantity based on an effective sample size of the data, itself potentially the product of a separate process of numerosity estimation.

Our results don’t allow us to tell with a high level of confidence whether the same weighting happens for the complexity penalty. Formally, the model that best describes our data is the “free scaling” BIC, where the complexity penalty scales logarithmically in the effective number of data-points *Ñ*. However, this model is closely followed in predictive power by the “free scaling” AIC, where the complexity penalty does not depend on *Ñ* at all (only the goodness-of-fit does). To some extent, it is unsurprising that it is harder to assess the dependence of the penalty (that is, to differentiate AIC from BIC) than that of the goodness-of-fit (differentiating, for instance, the distinct variants of AIC), because the logarithmic dependence of the BIC complexity on *Ñ* compresses the range over which it varies, and consequently its sensitivity to the specific values taken by *Ñ*. However, it is clear from all of our penalty-incorporating descriptions that the weight of the penalty is higher than what would be expected from the normative theory. In agreement with previous findings on a different task [1], our participants are therefore more skeptical of complex models than what would be recommended by an optimal strategy.

The main goal in our experimental design was to allow for a quantitative characterization of the way in which sample size affects intuitive model selection. This entailed some necessary limitations for the present study. First, we did not conduct an independent assessment of numerosity perception in our setup, or attempt to manipulate our participants’ sense of number by altering the way in which the stimuli were presented. Such manipulation would yield a strong test of the role of numerosity estimation in model selection, and is a promising direction for future research. However, an experiment addressing this issue would be impractical to perform with the range of models and sample sizes that we adopted in the present work. Second, our experiment uses simple, abstract one-dimensional stimuli, focusing exclusively on model dimensionality as complexity penalty. As shown in [1], more nuanced model features related to complexity can have a non-negligible effect on human choices. We don’t know what form of sample size scaling, if any, might affect these features. It also remains an open question whether the specific sublinear scaling we observed applies to different or more naturalistic choices of noisy data, and more generally to other sensory modalities. Third, our task was limited to low-dimensional models (1 ≤ *K* ≤ 5), motivated by the fact that many interesting perceptual and cognitive phenomena can be effectively conceptualized and studied empirically using relatively simple parametric descriptions of the sensory data [37–43]. While we expect that the qualitative features of our conclusions (the amount of available data affects intuitive model selection; its impact follows a sublinear trend) to have broad validity, and their quantitative details to generalize to tasks involving models of moderate dimensionality, making quantitative predictions for settings involving very high dimensional models (such as those commonly used in machine learning) is a harder task. In fact, the theoretical frameworks needed to characterize the complexity of very high-dimensional models may require a more powerful set of analytical tools than the ones we incorporated in our analyses (see for instance [44–46]). Establishing whether this regime is relevant for intuitive model selection, and how to tackle it with our approach, would be a natural next step.

A different type of limitation is that, while our data supports a heuristic-based origin for the logarithmic scaling, it cannot entirely rule out an alternative “computation” description. It could be argued that participants may actually carry out an approximate form of Bayesian integration, where the likelihood is computed on a subset of the available data-points obtained through a sub-sampling process, possibly guided by attentional or processing bottlenecks. If the size of this subset scales logarithmically with the total sample size, it could produce (depending on which points are sampled) the same sublinear growth in the goodness-of-fit term we observed. In this view, numerosity perception and likelihood calculation would not be feeding into each other, but would rather both be outcomes of the same underlying attentional sampling mechanism, thus displaying similar scaling. This interpretation, while conceivable, posits a complex integration process on top of a specific, arbitrarily-defined sampling step, and is therefore less parsimonious than the one we choose to put forward. The heuristic description we used, where a global perceptual estimate directly modulates the trade-off, provides a simpler and more direct cognitive explanation that also naturally explains the scaling shift observed at high *N*.

In conclusion, this study provides the first quantitative characterization of how the human simplicity bias scales with the amount of available evidence. The sublinear scaling we found represents a qualitative departure from the linear scaling prescribed by normative statistical frameworks such as AIC and BIC. Our results show that this sublinear trend is not arbitrary but, at least for small-to-moderate sample sizes, follows a logarithmic compression as one may expect in a numerosity perception setting. We therefore conclude that intuitive model selection is not guided by a formal statistical computation — not even in an approximate sense — but by a resource-rational heuristic that repurposes the brain’s native perceptual machinery to solve abstract inferential problems.

## 4 Methods

### 4.1 Behavioral experiment

In our preregistered behavioral task, participants were shown a certain number *N* of white vertical lines with varying horizontal positions on a gray background. As a visual aid, each line was paired with a small circle, in order to help the participants discern overlapping lines: the horizontal position of the circle corresponding to that of the paired line, while the vertical position was a random coordinate on the line sampled in such a way as to not overlap with the circles of other nearby lines (Fig. 2a, top).

On each trial, the horizontal position *x* ∈ [− 1, 1] of the vertical lines was generated according to the following rules:

1. A random number *K* ∈ [1, 5] was sampled: this was the number of latent sources that the data would be generated from;
2. The position *µ*_*i*_ of each of the *K* latent sources was sampled from a uniform distribution in [−1, 1];
3. The position of each vertical line was sampled from the resulting gaussian mixture distribution 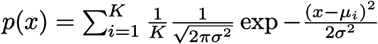, where *σ* = 0.1 was fixed (Fig. 2a, bottom).

Therefore, on each trial there were *K* clusters of vertical lines, sometimes with a certain degree of overlap between clusters. The participants had to report the number of latent sources, i.e., identify the model generating the data. They did so by pressing on their keyboard “W” to report 1, “D” to report 2, “K” to report 3, “O” to report 4, and Space to report 5. The key-to-number mapping was designed to allow a comfortable positioning of the hands on the keyboard (assuming a standard QWERTY layout) and hence avoid biases due to uncomfortably placed keys. To avoid confusion, this mapping was constantly indicated on the screen, the number selected by the participant was highlighted upon selection, and participants were allowed to correct mistakes by going back to the previous trial at any moment by pressing “B”.

The participants were instructed, during a brief introductory tutorial, on the properties of the Gaussian mixture models we used. In particular, they were shown different examples of data generated by the GMMs to illustrate the fact that each component had the same variance (and thus would generate clusters of approximately fixed width) and that components of the same model had the same weight (and thus would generate clusters of approximately the same size). To better help them internalize these concepts, participants were told that:

- the vertical lines were trails left on the ground by *N* wolves belonging to different packs: each area (i.e., each trial shown) could host from one to five packs;
- wolves tend to walk roughly in a file, so while walking each pack would leave a trail of approximately the same width, independently of the number of wolves in the pack;
- if more than one pack roamed an area, the sizes of the packs would approximately be the same.

Each participant was tested on 10 different values of *N* (number of vertical lines displayed): 16, 22, 28, 34, 40, 46, 52, 58, 64, and 70. For each value of *N*, participants were shown 90 trials. Unbeknownst to the participants, each trial was repeated 3 times in different moments of the experiment, in order to account for decision noise and verify participant consistency. Therefore, there were 30 distinct trials for each value of *N*, equally divided among the 5 values of *K*, for a total of 300 trials repeated 3 times each.

The trials were pre-generated with custom code and stored in a JSON-formatted data file. The file contained 30000 trials, 3000 for each *N*, of which 600 for each *K*. At the beginning of the experiment, 300 trials are selected at random from the file: 30 trials for each *N*, of which 6 for each *K*.

The task was divided into 30 small blocks of 30 trials each. In each block, all distinct trials for a given value of *N* were shown, in random order. The first block contained all trials with *N* = 70, the second block those with *N* = 64, and so on, in a progressively decreasing sequence of *N* until the last block, which contained the trials with *N* = 16. This sequence of blocks with decreasing values of *N* was then repeated 3 times. The participants were not informed of this repetition, and the trials were shuffled between blocks. The task was self-paced, and the participants were free to take breaks as they saw fit. Following the introductory tutorial, the participants performed a small number (60) of training trials. After each training trial, the participant was told whether their answer was correct. If it was not correct, we provided an assessment of how likely their answer was (*likely, fairly likely, unlikely, very unlikely*), according to the mathematical model used to generate the data for different values of *K*. This evaluation was based on the likelihood of the vertical positions of the lines according to a model with *K* components, where *K* corresponded to the answer given, and *µ*_*i*_ of each component was chosen to maximize the likelihood.

After the training phase, no more feedback was given to the participants on their performance. We ran the experiment on the online platform Pavlovia (pavlovia.org), with participants recruited on Prolific (prolific.com). We collected data from at least 100 participants who passed a pre-established performance threshold (25% or higher on each of the three sequences of blocks of the experiment; this threshold and the number of participants were fixed at preregistration). We discarded the data collected from all other participants. The final dataset included 100 participants (47% female; mean age = 34.16, std = 10.78, range 19-69). The average time required to complete the task was 75 minutes.

### 4.2 Experimental data analysis

To describe the behavior of our participants, we assumed that each of them sampled their choice for each trial from a probability distribution determined by a measure of goodness-of-fit *G* and a complexity penalty *P* for the models considered. We defined different sets of these terms, each connected to a possible explanation for the participants’ decision strategies. All the resulting probability distributions shared the same structure:

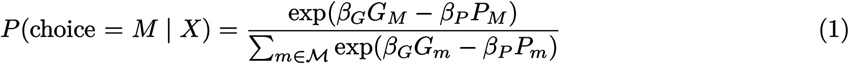

Where *β*_*G*_ and *β*_*P*_ represent how sensitive the participant is to the goodness-of-fit or the complexity penalty of the models, respectively, and ℳ is the set of five models the participants could choose from, as described in the previous section.

We used hierarchical Bayesian multinomial logistic regression to measure the parameters of each of the descriptions obtained from the different terms, and compared them using LOO to find the one fitting human behavior better. Here we explain each of the descriptions and the relative terms used, the details of the regressions, and of the comparison. All the descriptions, except for the “independent” one, were preregistered. The “independent” description was added upon realizing that it could be a reasonable and parsimonious candidate to describe human behavior, despite the scaling trend of the average answer observed in Fig 2c.

#### Log-likelihood description

To represent the hypothesis that participants lacked a simplicity bias, and only used a goodness-of-fit measure, we defined a model selection probability that only depends on the maximum log-likelihood *L*_*M*_ of each model for the trial data. The maximum log-likelihood for the Gaussian mixture models presented to the participants is given by:

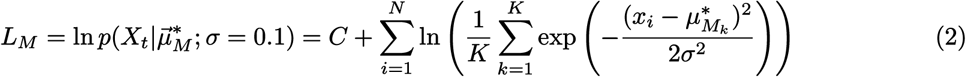

Where *X*_*t*_ is the data of trial *t, x*_*i*_ is the horizontal position of a singular data-point, *N* is the number of data-points, *K* is the number of components of the gaussian mixture distribution, 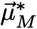 is the vector of component means that maximizes the likelihood for model *M* and 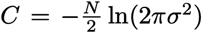 is a modelindependent constant. The resulting probability to choose the model *M* is:

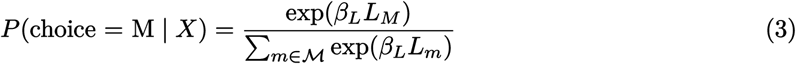

To find the values of *µ*^∗^ of each trial for each model, we used a modified Expectation-Maximization algorithm (EM algorithm, [47]). Since the standard deviation and weight of the components were fixed, to find the means we followed these steps:

1. Generate an initialization of 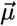;
2. Compute the *responsibility γ*_*nk*_ of each component *k* for each data-point *x*_*n*_:

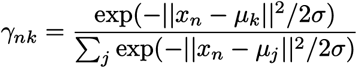
3. Compute each *µ*_*k*_ as the mean of the data-points weighted by the responsibility of *k*:

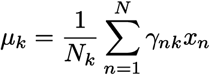
4. Repeat from step 2. until convergence.

Since the EM algorithm is only guaranteed to converge to a local maximum of the likelihood, we repeated the process with many different initializations, including:

- The positions of the true means that generated the data if *K* = *K*_true_;
- Every subset of size *K* of the original means if *K* < *K*_true_;
- The union of the true means and every possible subset of size *K* − *K*_*true*_ of the true means if *K* > *K*_true_. The means in the subsets were given a small displacement to avoid degeneracies;
- The centroids of a K-means clustering [48] with *K* components.

#### Cluster width and imbalance description

To represent the hypothesis that the log-likelihood doesn’t accurately capture how humans measure a model’s goodness-of-fit, and more generally that a different approach might better describe human behavior, we defined a model selection probability that depends on heuristics tailored to the stimuli. Each trial consisted of vertical lines grouped in clusters, generated by gaussian components with equal standard deviation and weight. Therefore, a natural way to determine if a K-component model is a good fit for a trial is to check if it produces clusters with a similar number of points and similar widths. More specifically, we considered the maximum width of the cluster (W) and the maximum imbalance in the number of points between clusters (I). These measures don’t penalize the complexity of a model, but only tell us how close the model’s partition of the data is to an optimal partition, according to the properties of Gaussian Mixtures. Therefore, similarly to the previous framework, this is a goodness-of-fit-based description.

To compute the terms, we clustered the data of each trial using the same EM algorithm described for the Log-likelihood description. This algorithm allows for overlapping clusters and favors balanced clusters when possible. The maximum width *W*_*M*_ of a model was defined as the maximum weighted standard deviation among all its clusters. The weighted standard deviation of each cluster was calculated using the responsibilities 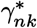 corresponding to the maximum likelihood means *µ*^∗^ of the model. Therefore, data-points with a higher probability of belonging to a cluster contribute more to its size calculation.

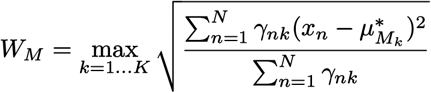

The maximum imbalance of points *I*_*M*_ is a measure of how evenly the data-points are distributed among its clusters. It was defined as the maximum size of any cluster compared to the optimal balanced size *N/K*. A perfectly balanced model, where all clusters are of equal size, will have a balance value of 1. A value greater than 1 indicates an imbalance, with some clusters being disproportionately more populated than others. The balance measure was computed by summing the responsibilities *γ*_*nk*_ of each cluster to determine its effective number of points, and then finding the largest of these normalized and scaled cluster sizes:

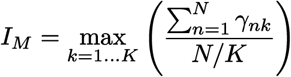

The resulting probability to choose model *M* is:

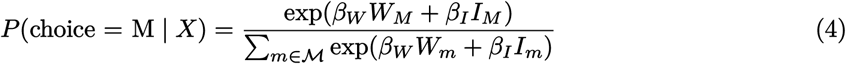

#### AIC and BIC “computation” description

We defined two descriptions of behavior based on two well-known Model Selection frameworks, the Akaike Information Criterion (AIC) and the Bayesian Information Criterion (BIC). These criteria are approximations of difficult-to-compute quantities. We used them as both a baseline to represent the hypothesis that participants display a simplicity bias in their strategies, and more specifically, to represent the hypothesis that strategies were based on a computation that parallels the ones used in theoretical model selection.

In the case of BIC, the computation being approximated is a Bayesian integration. BIC is an approximation of Bayesian Model Selection (BMS), based on the definition of “best model” as the one among a set with the highest posterior probability:

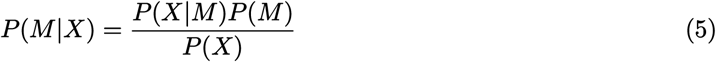

In our experiment, the model prior is uniform, so the only quantities needed to find the highest posterior probability are the marginal likelihoods of the data for each model:

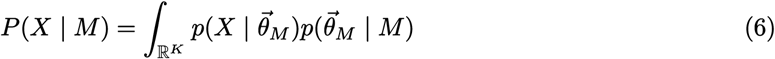

where 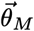 is the *K*-dimensional vector of model parameters. Using the Laplace approximation on the integral yields:

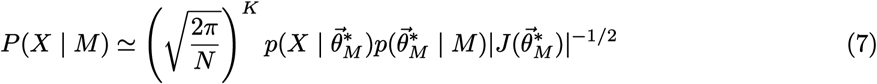

Applying the logarithm, and ignoring the *O*(1) terms, gives us BIC (Fig. 1a):

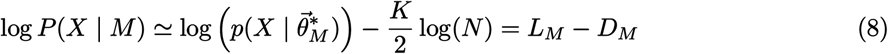

Therefore, if participants base their choices on Bayesian integration, or on an approximation of this computation, we expect their answers to display a scaling in *N* similar to that of a BIC-like description.

In the case of AIC, the criterion is derived from an estimate of the Kullback-Leibler divergence between the best fitting distribution 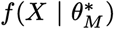 belonging to a parametric model *M* and the true distribution *g*(*X*) that generated the data:

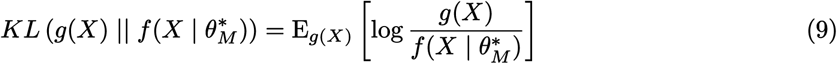

Where E_*g*(*X*)_ is the expected value under the distribution *g*(*X*). Using the properties of the logarithm, we can rewrite the divergence as:

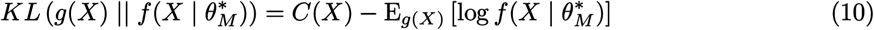

Where *C*(*X*) is a quantity that only depends on the data: when comparing two models, this quantity is the same for both of them, thus we can ignore it. The remainder 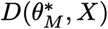 is still dependent on the true distribution *g*(*X*), a quantity that is usually inaccessible during model comparison. Akaike showed that under some assumptions, 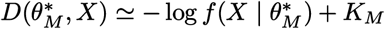, where *K*_*M*_ is the number of parameters of the model. Therefore, finding the model that minimizes the KL divergence corresponds to finding the one that maximizes *L*_*M*_ − *K*_*M*_.

While not directly derived from a computation that could be intuitively performed by participants, AIC finds the model with the maximum expected out-of-sample likelihood, i.e., the model that, on average, better describes unseen data. This makes it so that AIC approximates more computationally intensive methods, such as Cross Validation [49, 50]. Therefore, if participants base their choices on a computation that tries to find the highest predictive power among models, we expect their answers to display a scaling in *N* similar to that of an AIC-like description.

To compute the terms of both AIC and BIC “computation” descriptions, we used the same calculation of *L*_*M*_ presented for the Log-likelihood description. The penalty terms are trivial to compute. The resulting probabilities to choose model *M* are:

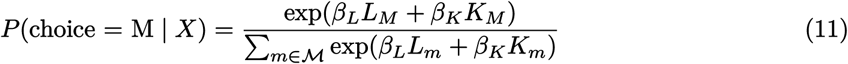

for AIC, and

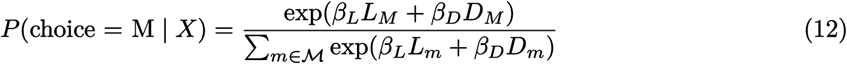

for BIC.

#### AIC “independent” description

To represent the hypothesis that participants lack any scaling, we defined a description of their behavior that doesn’t depend on the number of data-points *N*. On average, the log-likelihood grows linearly in the number of data-points. Indeed

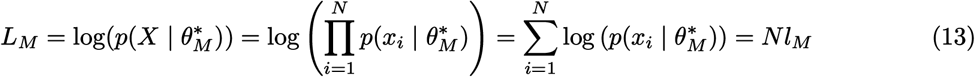

Where 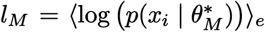, where ⟨·⟩_*e*_ is the empirical mean. This quantity also corresponds to the negative of the Kullback-Leibler divergence between the empirical measure of the data 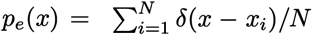 (where *δ*(*x* − *x*_*i*_) is Dirac’s delta) and the candidate model distribution, up to a model-independent additive value:

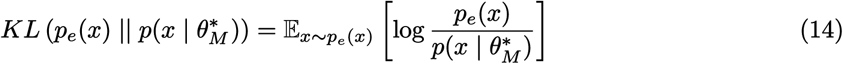

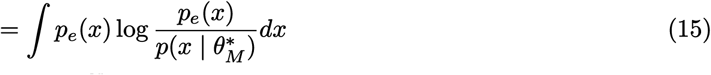

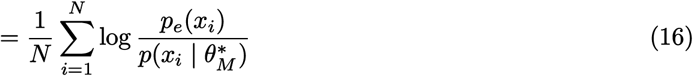

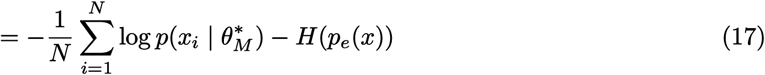

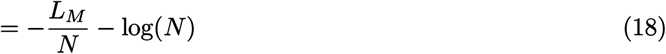

Where *H*(*p*_*e*_(*x*)) = log(*N*) is the entropy of the empirical measure.

We used *l*_*M*_ as the goodness-of-fit term of this description. For the penalty, we used the complexity penalty of AIC, *K*_*M*_, as it is independent of *N*. The disproportion in relative size of goodness-of-fit and penalty after these adjustments is balanced in the regression by the *β* sensitivities, which will therefore result bigger for the goodness-of-fit in this description compared to the previous ones.

The resulting probability to choose model *M* is:

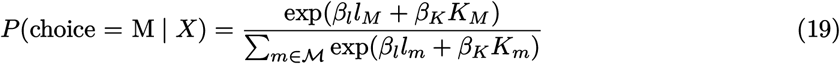

#### AIC and BIC “heuristic scaling” descriptions

To represent the hypothesis that participants adopt strategies based on a heuristic rebalancing of the relative weight of goodness-of-fit and complexity penalty, based on on an “effective” perceived size of the data, we defined two descriptions where the terms depend not on *N*, but on *Ñ* = log(*N*). The choice for a sublinear function was driven by the literature on numerosity perception, where a compression on the perceived quantity of data is well documented [23, 25, 28, 51]. We specifically chose the logarithm function as a compressive function to parallel Weber’s law [27], which has been shown to govern numerosity perception in humans [24, 26].

The goodness-of-fit term with logarithmic scaling was defined as *Ñ* · *l*_*M*_ = log(*N*) · *l*_*M*_, where log is the natural logarithm. For the AIC-based description, the complexity penalty remained unchanged. For the BIC-based description, it was defined as 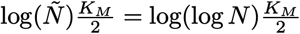. Since the values of *N* we used were all greater than Euler’s number *e*, there were no undefined or negative values of log(*Ñ*).

The resulting probabilities to choose model *M* are:

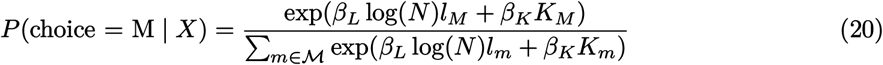

for AIC and

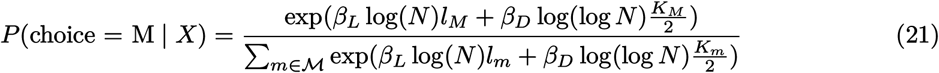

for BIC.

#### AIC and BIC “free scaling” descriptions

To find the actual shape of the scaling of our participants, we defined a modified version of the “heuristic scaling” descriptions. In these versions, *Ñ* was not defined as the logarithm of *N*, but as a free function to be inferred. For simplicity, we assumed this function to be the same for all participants. The details of the inference are presented below.

The resulting probabilities to choose model *M* are:

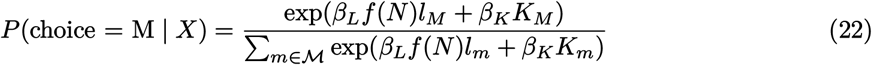

for AIC and

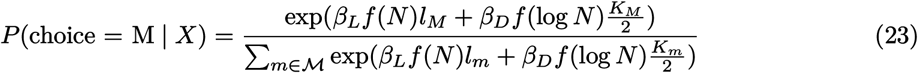

for BIC.

#### Lapse rate

We extended all the defined probabilities to account for lapses by defining a lapse rate *ϵ* for each participant. Therefore, on each trial, there is a probability (1 − *ϵ*) that the participant will perform the task and act following their strategy, and a probability *ϵ* that they will answer randomly. The general structure of the probability to choose model *M* will therefore be:

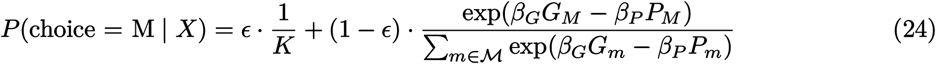

Where 1*/K* represents the uniform probability to randomly choose model *M*.

To be able to perform model comparison reliably, we also added a second lapse rate *δ* (see Model comparison). While *ϵ* is the lapse rate of a single participant, the same *δ* is in common between all participants. Philosophically, this term represents the subject-independent, global probability that external events will lead to errors in the execution of the task. From a practical point of view, this term can be included in *ϵ* by slightly changing its definition: if we define as 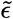 the lapse of a participant, their overall lapse will become 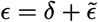. Therefore, adding *δ* doesn’t affect the above formula, but only alters the prior distribution of *ϵ*.

### 4.3 Bayesian regression

In the formulas for each description of possible strategies we defined, the probability to choose a model *M, P*(M | *X*), depends on the subject-specific parameters 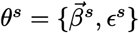 (where 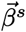 represents the parameters of a specific strategy for subject *s*). To infer their values we used a hierarchical Bayesian approach [30]. We used the same regression structure for all strategies other than the “free scaling” ones, which are described in more detail in the next section.

For all the descriptions, denoting by *C* the recorded choices of all participants, and by 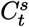 the recorded choice of participant *s* on trial *t*, we define the likelihood function of the associated parameters *θ*^*s*^ is:

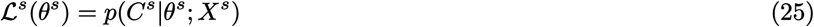

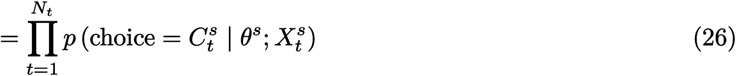

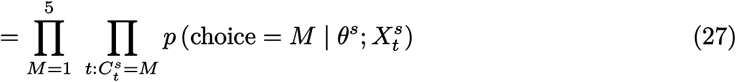

We also define a prior for each of the parameters:

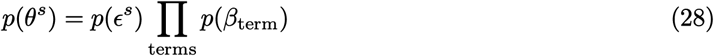

where the terms are specific to each description.

By applying Bayes’ theorem, we compute a joint posterior probability distribution for the parameters:

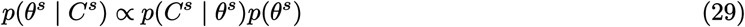

To extend the same computation to the entire population we use a hierarchical approach. This allows us to infer population-level parameters and, at the same time, regularize the inference procedure, with a regularization strength that depends on the data [30]: since the participants belong to the same population and are exposed to the same experimental conditions, we assume that the parameters describing them will be related. In other words, the data we collect about one participant tells us something about that specific participant, but should also inform us about human behavior in general, and thus about the behavior of other participants. We can build this assumption into the model by imposing that the parameters associated with different participants are all sampled from the same distribution, leaving the parameters of this higher-level probability distribution to be also inferred from the data. This procedure has the additional advantage of partially constraining the subject-level parameters to reasonable values given what we observed in the other participants, thus acting as a regularizer for the inference process. Therefore, our posterior will be:

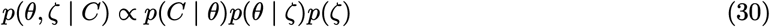

where 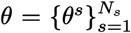 are the subject-specific parameters for all *N*_*s*_ participants, and *ζ* are the common (population) parameters.

To ensure a more robust inference, the terms of each description (corresponding to the regression predictors) have been standardized to a mean of zero and a standard deviation of one before fitting the model. The “free scaling” descriptions’ terms, as previously stated, could not be standardized, since they are computed during the inference process. Therefore, we standardized *L*_*M*_ before multiplying it by *f*(*N*)/*N*. For the dimensionality term, we standardized *K*_*M*_ /2 before multiplying it by log(*f*(*N*)). In the following, we will refer to the parameters as 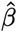 when they are associated with a standardized term and *β* when they are not.

In the likelihood-only description, we also added to each likelihood, before standardization, a small noise ∈ [− 0.5, 0.5] · 10^−5^: this small shift ensured a stable inference in those trials where more than one model had the exact same likelihood. Indeed, for those trials, the probabilities assigned by the models (*p* ∝ exp(*β*_*L*_*L*_*M*_) = 0.2 if all the models have the same likelihood) are independent of *β*_*L*_. This caused the same importance weights to be assigned to all the parameters of the MCMC chain for these trials (see Model comparison), which in turn made the Pareto inference of the PSIS LOO computation unreliable.

The specific priors we assign to the parameters (*p*(*θ* | *ζ*) in Eq (30)) are:

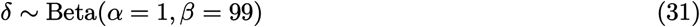

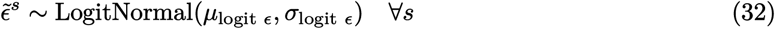

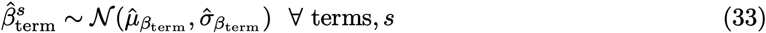

Our population parameters, *ζ*, are therefore the logit mean *µ*_logit *ϵ*_ and standard deviation *σ*_logit *ϵ*_ of the 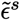 parameters, and 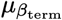 and 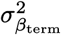 for the *β*^*s*^ parameters of each term. Following standard practice, we assign weakly informative priors *p*(*ζ*) to the population-level parameters:

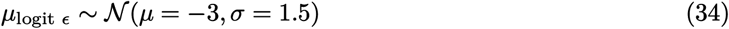

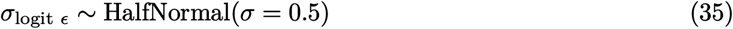

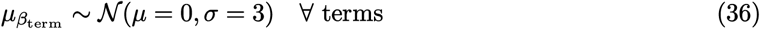

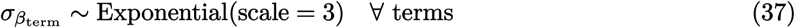

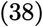

The same priors were assigned to all descriptions, with the only exception of the Log-likelihood description, which had a prior

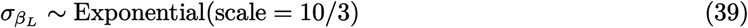

for its only *β* term. In the definition of each description, these priors were implemented through the standardized coefficients and the empirical standard deviation of each term, thus defining the distributions of 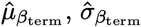, and 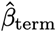.

Finally, the global likelihood function was obtained as a product of the subject-level likelihood functions:

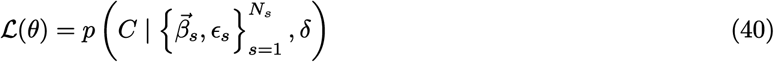

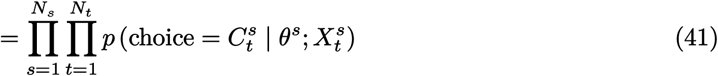

#### Gaussian Processes

In the “free scaling” descriptions, the scaling of both goodness-of-fit and penalty terms is a generic function *f*(*N*). To infer the shape of this function from the data, we employed a Gaussian Process (GP) [35]. GPs are a principled, non-parametric method for defining a prior over functions, allowing us to treat the whole function as a parameter of the regression.

The GP prior was defined as:

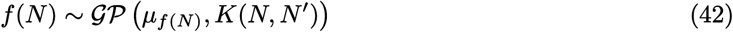

where *µ*_*f*(*N*)_ is the mean of the prior for each value of *N* (typically set to *µ*_*f*(*N*)_ = 0) and *K*(*N, N*^*′*^) is the Exponentiated Quadratic covariance kernel. A GP kernel is a function that allows us to compute the covariance between any two values of *N, N*_*i*_, and *N*_*j*_:

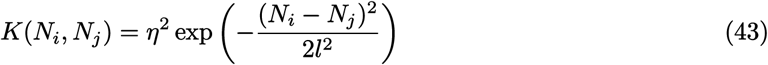

where *η* is the marginal standard deviation of the Gaussian process, which controls the overall vertical amplitude of the function, and *l* is the length scale, which determines its smoothness. We placed weakly informative priors on these hyperparameters:

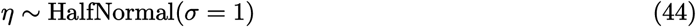

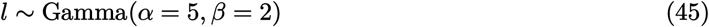

A challenge in this approach is potential model degeneracy: if both the regression coefficients (*β*_*L*_ or *β*_*D*_) and the function *f*(*N*) (modeling, together, the goodness-of-fit term and the dimensionality penalty) are flexible, the model can become unidentifiable. For instance, one can double the scale of the GP function while halving the *β*_*L*_ coefficient without changing the likelihood. To resolve this non-identifiability issue, we enforce the constraint *f*(*N* = 16) = 16. This is equivalent to using the GP function to model scaling in a relative sense, leaving the absolute magnitude of the goodness-of-fit and dimensionality terms to be controlled by the regression coefficients *β*.

To implement this constraint, we defined the GP as a conditional multivariate normal distribution. Given the full covariance matrix *K* over all unique values of *N*, we partition it as follows:

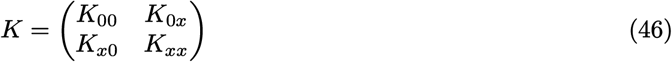

where *K*_00_ is the variance at the reference point (*N* = 16), *K*_*xx*_ is the covariance matrix of the other points, and *K*_*x*0_ contains the cross-covariances. The values of the GP for all values of *N* other than 16, *f*(*N*_*x*_), are then drawn from a multivariate normal distribution whose parameters are conditioned on the fixed value *f*(16) = 16:

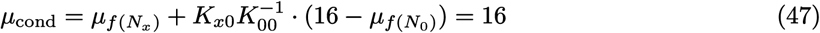

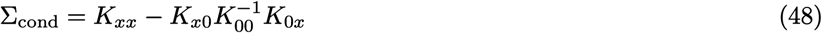

Where, to impose *f*(16) = 16, we put *µ*_*f*(*N*)_ = 16 for all *N*.

To ensure numerical stability and improve sampling efficiency, we adopt a non-centered parameterization. Instead of sampling directly from this correlated conditional distribution, we sampled a vector of independent standard normal variables *z* and transformed them using the Cholesky decomposition [52] of the conditional covariance matrix, *L* = chol(Σ_cond_), so *f*(*N*_*x*_) = *µ*_cond_ + *L* · *z*. Moreover, we rescaled the input values of the GP (i.e., of the kernel function) as 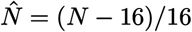.

#### Sampling the posterior distribution

We performed inference with a Markov Chain Monte Carlo (MCMC) method using the NUTS algorithm [53], as implemented in PyMC version 5.25.1 [54]. This method samples parameters from the posterior distribution *p*(*θ* | *C*): with a high enough number of samples, the distribution of the samples closely matches the distribution of the posterior. We sampled 4 independent chains of 10000 draws each, 5000 of which we used for tuning the algorithm and subsequently discarded, and 5000 as actual samples of the target posterior distribution. For computational convenience, we applied a thinning process to reduce the number of posterior samples, keeping only 1000 for the analyses. The target acceptance probability parameter for NUTS was set at 0.95, and no divergences were detected during sampling (see Tables 1 to 9).

### 4.4 Model comparison

To verify which of the descriptions better fitted human behavior, we compared them with a Leave-One-Out Cross-Validation (*LOO*) estimate of the Expected Log pointwise Predictive Density (*ELPD*). LOO estimates how well, on average, the model (i.e., a description) describes data it hasn’t been trained on. To do so, LOO computes how well the result of a regression on all the trials (across all participants) except one predicts the outcome of that trial, and averages this value over all the trials. The ELPD, therefore, automatically balances goodness of fit with model complexity, penalizing overly complex models that tend to overfit the noise in the data. More formally, the LOO estimate of the ELPD for a statistical model ℳ for which we can sample the posterior distribution with MCMC is defined as:

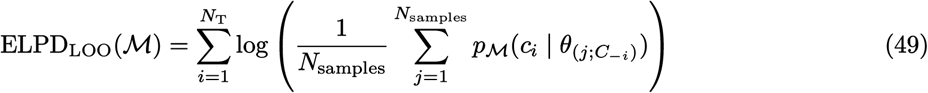

where *N*_T_ is the total number of trials in the data, *N*_samples_ is the number of posterior samples in the Markov chain for model ℳ, *p*_ℳ_ is the likelihood function of the model, *c*_*i*_ is the data (i.e., the human choice) from the *i*-th trial and 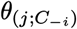 is the *j*-th posterior sample of a Markov chain generated from all data except *c*_*i*_ (note that each posterior distribution sample, including 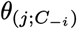, is a high-dimensional vector containing one value for each parameter in the model). In Eq (49), the argument of the logarithm represents the likelihood of the description ℳ for the choice *c*_*i*_, averaged over the posterior distribution over *θ* (where this posterior is obtained without looking at data-point *c*_*i*_). Indeed, since the training of ℳ results in a posterior distribution rather than a single parameter vector, all quantities related to ℳ must be averaged over this distribution. For MCMC training, this average is computed as a sum over all sampled parameters, given that the samples follow the posterior distribution: 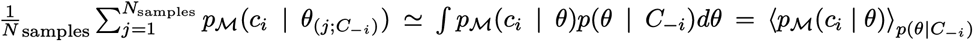. In practice, computing Eq 49 requires training the model *N*_*T*_ times, one per trial: given the large number of trials in our experiment, we approximated the leave-one-out procedure with Pareto-smoothed importance sampling (PSIS-LOO) [32].

Importance Sampling LOO is based on the fact that, under the hypothesis of conditional independence of the data given the parameters,

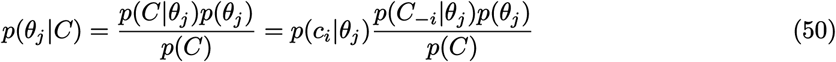

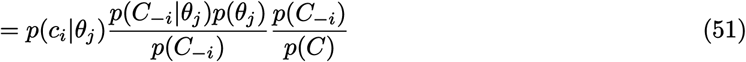

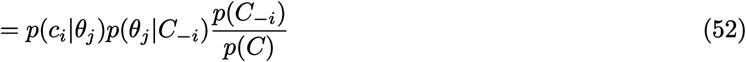

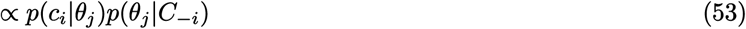

Therefore,

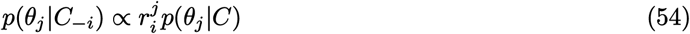

where

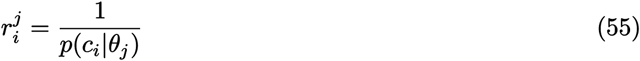

which implies that we can approximate a Monte Carlo average over *p*(*θ*_*j*_ |*C*_−*i*_) by one over *p*(*θ*_*j*_ | *C*), where each sample is weighted by the importance weight 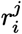. So instead of writing

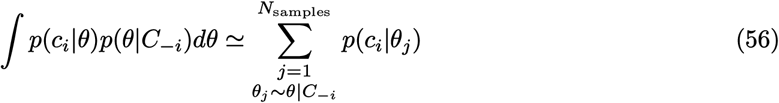

we can write

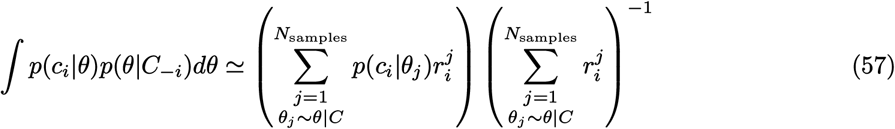

Using the 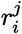 as weights, we can approximate the sum on *N*_*samples*_ in Eq. 49 as:

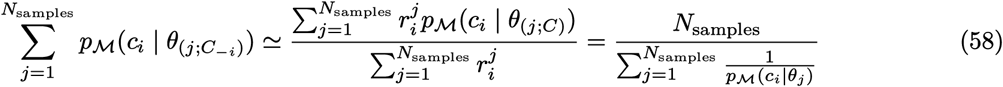

The problem with this approximation is that *p*_ℳ_ (*c*_*i*_ | *θ*_*j*_) can be arbitrarily small, leading to instabilities, as a single element of the chain can greatly skew the result. To avoid this, a Pareto distribution is fitted on the *M* biggest 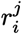 (corresponding to the smallest *p*_ℳ_ (*c*_*i*_ | *θ*_*j*_)) for each *i*, and the weights are substituted by new weights sampled from the fitted distribution, with a threshold to avoid sampling extremely high weights.

#### Necessity of *δ* in the lapse rate

In our dataset, a few participants lapsed (or were considered lapsing by the description) only one time. For the trained model, this means that the participant has *ϵ* > 0. However, during the computation of *LOO*, the exclusion of that trial results in *ϵ* ≃ 0. This means that the trial alone guides the value of the inferred *ϵ*: it’s a highly influential observation for the regression. These kinds of observations are not problematic for LOO, but can greatly affect the performance of the PSIS approximation. To avoid this kind of trial being too influential, we included the parameter *δ*: this parameter gives *ϵ* a lower bound, computed across all participants, thus avoiding the possibility that removing a single trial will influence the inferred lapse rate too much.

#### Necessity for the added noise to *L*_*M*_ for the likelihood-only description

For the likelihood-only description, for some trials all the probabilities *p*_ℳ_ (*c*_*i*_ | *θ*_*j*_) are the same, independently of *θ*_*j*_, for all *j*. This causes problems when fitting the Pareto distribution, as the weights are all equal and cannot be meaningfully interpreted as being Pareto distributed. The fitted Pareto distribution has an infinite variance, making the Pareto Smoothed Importance Sampling unreliable. To avoid this problem, we added a small noise to all the likelihoods, thus breaking this symmetry and introducing differences in the values of the weights.

## Data availability statement

The data collected in this study, the source code for model fitting and generating the figures, as well as the MCMC chains derived from the data and used to perform statistical analyses are available in a publicly accessible repository on Zenodo [55].

## 4.5 Supplementary Material

^1^In [1], the dimensionality is computed slightly differently as 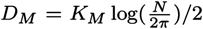. However, the over-weighting of dimensionality persists even when adjusting for this difference, i.e., when multiplying the sensitivity to dimensionality *β*_*D*_ estimated in [1] by 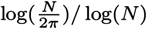.

**Supplementary Figure S1.**
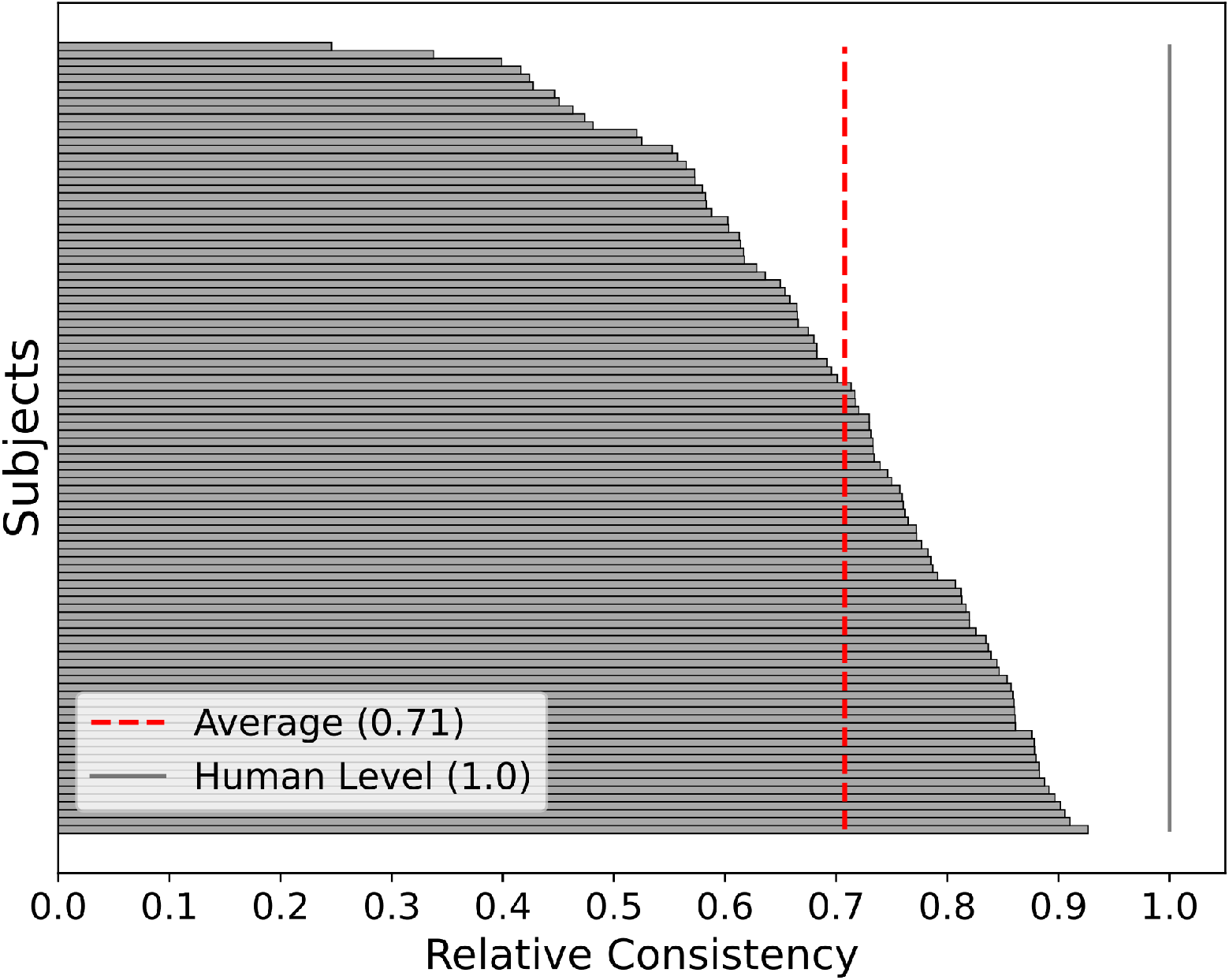
Relative description performance per participant. Normalized relative consistency for each individual subject, calculated by scaling the absolute description consistency between the random guessing chance (set to 0) and the participant’s own self-consistency (set to 1.0, marked by the vertical gray line). Horizontal bars represent individual participants, sorted in order of their relative consistency score. The vertical dashed red line indicates the average across all participants.

**Table 3:** Posterior summary statistics for the key population-level parameters of the AIC “independent” description. Details are as in Table 1.

|  | mean | sd | hdi_3% | hdi_97% | mcse_mean | mcse_sd | ess_bulk | ess_tail | r_hat |
| --- | --- | --- | --- | --- | --- | --- | --- | --- | --- |
| $\mu_{\beta_L}$ | 9.480 | 0.638 | 8.268 | 10.687 | 0.010 | 0.007 | 4087.1 | 4058.3 | 1.000 |
| $\mu_{\beta_D}$ | 1.482 | 0.064 | 1.367 | 1.607 | 0.001 | 0.001 | 3969.8 | 3970.8 | 1.001 |
| $\sigma_{\beta_L}$ | 6.301 | 0.489 | 5.428 | 7.248 | 0.008 | 0.005 | 4053.8 | 3774.1 | 1.001 |
| $\sigma_{\beta_D}$ | 0.629 | 0.047 | 0.541 | 0.712 | 0.001 | 0.001 | 3971.0 | 3969.8 | 1.000 |
| $\mu_\epsilon$ | 0.023 | 0.003 | 0.017 | 0.028 | 0.000 | 0.000 | 3465.2 | 3558.8 | 1.000 |

**Table 4:** Posterior summary statistics for the key population-level parameters of the AIC “heuristic scaling” description. Details are as in Table 1.

|  | mean | sd | hdi_3% | hdi_97% | mcse_mean | mcse_sd | ess_bulk | ess_tail | r_hat |
| --- | --- | --- | --- | --- | --- | --- | --- | --- | --- |
| $\mu_{\beta_L}$ | 2.712 | 0.177 | 2.384 | 3.052 | 0.003 | 0.002 | 4123.2 | 3595.0 | 1.000 |
| $\mu_{\beta_D}$ | 1.474 | 0.063 | 1.358 | 1.593 | 0.001 | 0.001 | 3895.7 | 3726.4 | 1.001 |
| $\sigma_{\beta_L}$ | 1.773 | 0.139 | 1.506 | 2.022 | 0.002 | 0.002 | 4028.6 | 3891.3 | 1.001 |
| $\sigma_{\beta_D}$ | 0.624 | 0.047 | 0.533 | 0.709 | 0.001 | 0.001 | 3721.5 | 3865.0 | 1.000 |
| $\mu_\epsilon$ | 0.023 | 0.003 | 0.017 | 0.028 | 0.000 | 0.000 | 3449.3 | 3562.5 | 1.000 |

**Table 5:** Posterior summary statistics for the key population-level parameters of the BIC “heuristic scaling” description. Details are as in Table 1.

|  | mean | sd | hdi_3% | hdi_97% | mcse_mean | mcse_sd | ess_bulk | ess_tail | r_hat |
| --- | --- | --- | --- | --- | --- | --- | --- | --- | --- |
| $\mu_{\beta_L}$ | 2.714 | 0.179 | 2.375 | 3.040 | 0.003 | 0.002 | 3942.8 | 3541.6 | 1.001 |
| $\mu_{\beta_D}$ | 0.813 | 0.036 | 0.743 | 0.877 | 0.001 | 0.000 | 3924.8 | 3710.5 | 1.000 |
| $\sigma_{\beta_L}$ | 1.768 | 0.139 | 1.510 | 2.032 | 0.002 | 0.002 | 3641.3 | 3506.6 | 1.002 |
| $\sigma_{\beta_D}$ | 0.354 | 0.027 | 0.305 | 0.404 | 0.000 | 0.000 | 3916.2 | 3651.8 | 1.000 |
| $\mu_\epsilon$ | 0.024 | 0.003 | 0.018 | 0.029 | 0.000 | 0.000 | 2843.0 | 3272.3 | 1.000 |

**Table 6:** Posterior summary statistics for the key population-level parameters of the AIC “free scaling” description. Details are as in Table 1.

|  | mean | sd | hdi_3% | hdi_97% | mcse_mean | mcse_sd | ess_bulk | ess_tail | r_hat |
| --- | --- | --- | --- | --- | --- | --- | --- | --- | --- |
| $\mu_{\beta_L}$ | 0.526 | 0.037 | 0.459 | 0.594 | 0.001 | 0.000 | 3999.2 | 3830.4 | 1.001 |
| $\mu_{\beta_D}$ | 1.480 | 0.064 | 1.357 | 1.594 | 0.001 | 0.001 | 4257.3 | 3936.6 | 1.000 |
| $\sigma_{\beta_L}$ | 0.342 | 0.027 | 0.291 | 0.393 | 0.000 | 0.000 | 4139.2 | 3861.4 | 1.003 |
| $\sigma_{\beta_D}$ | 0.627 | 0.047 | 0.539 | 0.714 | 0.001 | 0.001 | 4102.7 | 3879.6 | 1.000 |
| $\mu_\epsilon$ | 0.023 | 0.003 | 0.017 | 0.028 | 0.000 | 0.000 | 2729.2 | 3501.0 | 1.001 |
| $\eta_{pop}$ | 2.230 | 0.509 | 1.380 | 3.249 | 0.008 | 0.006 | 4297.2 | 3834.1 | 1.001 |
| $l_{pop}$ | 2.049 | 0.477 | 1.195 | 2.938 | 0.007 | 0.005 | 4073.7 | 3916.5 | 1.001 |

**Table 7:** Posterior summary statistics for the key population-level parameters of the BIC “free scaling” description. Details are as in Table 1.

|  | mean | sd | hdi_3% | hdi_97% | mcse_mean | mcse_sd | ess_bulk | ess_tail | r_hat |
| --- | --- | --- | --- | --- | --- | --- | --- | --- | --- |
| $\mu_{\beta_L}$ | 0.489 | 0.035 | 0.423 | 0.551 | 0.001 | 0.000 | 3437.3 | 3828.6 | 1.001 |
| $\mu_{\beta_D}$ | 0.984 | 0.042 | 0.896 | 1.057 | 0.001 | 0.001 | 3552.9 | 3633.2 | 1.002 |
| $\sigma_{\beta_L}$ | 0.318 | 0.027 | 0.271 | 0.371 | 0.000 | 0.000 | 3170.7 | 3647.7 | 1.001 |
| $\sigma_{\beta_D}$ | 0.419 | 0.032 | 0.365 | 0.482 | 0.001 | 0.000 | 4044.8 | 3827.8 | 1.000 |
| $\mu_\epsilon$ | 0.022 | 0.003 | 0.017 | 0.028 | 0.000 | 0.000 | 2809.1 | 3213.3 | 1.002 |
| $\eta_{pop}$ | 2.715 | 0.527 | 1.806 | 3.749 | 0.009 | 0.006 | 3803.9 | 3775.3 | 1.001 |
| $l_{pop}$ | 1.457 | 0.386 | 0.770 | 2.151 | 0.007 | 0.005 | 3264.1 | 3817.0 | 1.001 |

**Table 8:** Posterior summary statistics for the key population-level parameters of the L-only description. Details are as in Table 1.

|  | mean | sd | hdi_3% | hdi_97% | mcse_mean | mcse_sd | ess_bulk | ess_tail | r_hat |
| --- | --- | --- | --- | --- | --- | --- | --- | --- | --- |
| $\mu_{\beta_L}$ | 0.057 | 0.006 | 0.046 | 0.067 | 0.000 | 0.000 | 4140.3 | 3592.9 | 1.000 |
| $\sigma_{\beta_L}$ | 0.057 | 0.005 | 0.048 | 0.066 | 0.000 | 0.000 | 4129.7 | 3805.3 | 1.000 |
| $\mu_\epsilon$ | 0.014 | 0.002 | 0.011 | 0.019 | 0.000 | 0.000 | 3949.8 | 4058.4 | 1.000 |

**Table 9:** Posterior summary statistics for the key population-level parameters of the W-I description. Details are as in Table 1.

|  | mean | sd | hdi_3% | hdi_97% | mcse_mean | mcse_sd | ess_bulk | ess_tail | r_hat |
| --- | --- | --- | --- | --- | --- | --- | --- | --- | --- |
| $\mu_{\beta_W}$ | -12.488 | 0.825 | -14.098 | -11.001 | 0.013 | 0.009 | 3948.2 | 3615.3 | 1.000 |
| $\mu_{\beta_I}$ | 0.201 | 0.109 | -0.003 | 0.406 | 0.002 | 0.001 | 4075.8 | 3918.7 | 1.000 |
| $\sigma_{\beta_W}$ | 8.209 | 0.648 | 6.987 | 9.391 | 0.010 | 0.007 | 3914.4 | 3854.9 | 1.001 |
| $\sigma_{\beta_I}$ | 1.075 | 0.083 | 0.927 | 1.234 | 0.001 | 0.001 | 3905.7 | 3439.7 | 1.000 |
| $\mu_\epsilon$ | 0.004 | 0.001 | 0.002 | 0.006 | 0.000 | 0.000 | 3581.5 | 3764.7 | 1.001 |

